# Linking brain structure to stress reactivity: Cingulate surface area predicts acute cortisol responses

**DOI:** 10.1101/2025.09.12.675570

**Authors:** Emin Serin, Lea Sophie Schill, Christoph Bärtl, Marina Giglberger, Julian Konzok, Hannah L. Peter, Nina Speicher, Ludwig Kreuzpointner, Brigitte M. Kudielka, Stefan Wüst, Henrik Walter, Gina-Isabelle Henze

## Abstract

**Background:** Altered stress responses are closely linked to mental disorders, but the role of brain structure in acute cortisol responses to psychosocial stress remains underexplored, particularly in healthy individuals. Previous studies, with predominantly small samples, primarily focused on selected limbic regions and functional measures. Thus, this study investigates associations between brain structure and cortisol responses to psychosocial stress, exploring if hypothalamic-pituitary-adrenal axis reactivity can be predicted from brain morphology.

**Methods:** Our study included 291 subjects (157 females, 18-62 years) and consisted of two parts. First, a confirmatory analysis examined associations between specific cortical surface area, thickness, and subcortical volume with stress-induced cortisol increases using Permutation Analysis of Linear Models (PALM). Second, we conducted an exploratory whole-brain vertex-wise analysis, followed by out-of-sample prediction of cortisol increases from structural measures.

**Results:** We found consistent negative associations between cingulate cortex (CC) sub- structures and acute cortisol increases. In PALM- and whole-brain analysis, a smaller surface area of the left rostral and caudal anterior cingulate cortex (cACC), posterior cingulate cortex, and right cACC were associated with higher cortisol stress responses, particularly in males. The left cACC surface area emerged as the most promising predictor in machine learning analyses. Additionally, other fronto-limbic structures were also associated with or predictive of acute cortisol reactivity.

**Conclusions:** Our findings demonstrate that cortical and subcortical structural measures, particularly smaller surface areas of the CC, predict acute hormonal stress responses. Notably, the left cACC emerged as the most consistent predictor, underlining its potential as a biomarker for stress-related diseases.

## INTRODUCTION

Stress is an integral part of everyday live and the confrontation with actual or perceived stressors disrupts the body’s homeostasis (Sapolsky, 2003), triggering both physiological and psychological stress responses (Herman et al., 2003). Acute and chronic stress have been linked to a range of negative effects on physical and mental health (Chrousos, 2009). Research conducted on animals and humans suggests that these harmful effects may stem, at least partly, from prolonged exposure to glucocorticoids such as corticosterone and cortisol (Lupien et al., 1998; McEwen, 2002). Consequently, neural mechanisms underlying the (acute) cortisol stress response have gained much attention in human neuroscientific research in recent years. In this context, several psychosocial stress induction paradigms have been developed for laboratory use as well as imaging environments (functional magnetic resonance imaging, fMRI) and have been shown to reliably elicit increases in cortisol levels (Berretz et al., 2021; Dickerson & Kemeny, 2004; Noack et al., 2019; Qiu et al., 2022). Most studies have focused on Blood-Oxygenation-Level-Dependent (BOLD) measures, such as resting state functional connectivity (rsFC) and task-based responses, though the exact nature of the reported effect patterns remains debated (Harrewijn et al., 2020; Henze et al., 2020, 2021; Noack et al., 2019; Van Oort et al., 2017). Although structural, i.e. T1- and/or T2-weighted (T1_w_/T2_w_-) images are a by-product of such neuroimaging studies, few analyses have explored associations between structural measures of the brain and acute cortisol stress responses in healthy subjects (Henze et al., 2023). Given that numerous studies indicate aberrant brain structural parameters in the context of stress-related disorders (Cardoner et al., 2024; Kaul et al., 2021; McEwen et al., 2016; Radley & Morrison, 2005), it is surprising that the relationship between brain structure and stress-induced cortisol release in a healthy organism remains poorly understood.

Most studies on stress-induced changes in brain structure focus on the hippocampus due to its sensitivity to chronic stress and glucocorticoids, as well as its role in learning, memory, and hypothalamic-pituitary-adrenal (HPA) axis regulation (Herman et al., 2003, 2005, 2016; Jankord & Herman, 2008). Moreover, prolonged glucocorticoid-exposure, e.g., due to chronic stress, have been linked to hippocampal volume reduction and neurotoxic effects on neurons involved in learning and memory (Cardoner et al., 2024; Kaul et al., 2021; McEwen et al., 2016; Radley & Morrison, 2005). Other regions, such as the medial prefrontal cortex (mPFC) and amygdala, have gained attention for their importance in orchestrating stress responses and the processing, acquisition, and extinction of fear memories (Alexandra Kredlow et al., 2022; Herman et al., 2016; Jankord & Herman, 2008; Van Oort et al., 2017). Thus, volume and thickness reductions in these areas appear to be more likely depending on the stress-related condition (e.g., post-traumatic stress disorder, PTSD; major depressive, and anxiety disorder; 16,17). The few studies reporting on associations between brain structure and acute cortisol stress reactions in response to psychosocial stressors in healthy human subjects also focused on these regions (Henze et al., 2023). However, it is becoming increasingly clear that other striato- limbic structures also deserve significant attention (Cardoner et al., 2024; Henze et al., 2023; Kaul et al., 2021) and that examining surface area may provide additional insights (Fowler & Gaffrey, 2022; Hu et al., 2018; Panizzon et al., 2009; Rimol et al., 2012; Stomby et al., 2016; Sun et al., 2019).

To our knowledge, twelve studies have applied psychosocial stressors in humans to elicit acute cortisol responses and studied their associations with structural brain measures. Of these, only two studies conducted a whole-brain analysis (J. Liu et al., 2012; Uhlig et al., 2023), while the rest carried out region of interest (ROI-) analyses on two, hippocampus and amygdala (Admon et al., 2017; Fowler & Gaffrey, 2022), or even just one ROI: hippocampus (Blankenship et al., 2019; Degering et al., 2023; J. C. Pruessner et al., 2008; M. Pruessner et al., 2007), amygdala (Barry et al., 2017; Klimes-Dougan et al., 2014), and perigenual ACC (pACC, (Boehringer et al., 2015). Only one study examined more than two ROIs and thereby considered both subcortical volume and cortical thickness (Henze et al., 2023). However, no consistent pattern of association emerges from these studies, which is probably due to several factors: Variations in focus on different aspects of the cortisol response (such as total cortisol release, acute increase, recovery phase, or distinct response types), small sample sizes (average *n* = 54), and unaccounted factors like age, sex, gender, or, in females, menstrual cycle phase, hormonal contraception, and reproductive status. Additionally, differences in stress induction paradigms (e.g., Trier Social Stress Test or similar methods) and methodological inconsistencies (such as timing of stress induction or assessing cortisol and structural data on different days) further contributed to the discrepancies.

In a recent study (Henze et al., 2023), we have addressed some of these aspects using Permutation Analysis of Linear Models (PALM), controlling for age, sex, and total brain volume (TBV), to investigate the relationship between (sex-specific) cortisol increases to acute psychosocial stress and anatomical variables of twelve striato-limbic structures (i.e., volume and thickness, lateralized). Scan*STRESS*, a psychosocial stress induction paradigm for imaging environments based on the TSST (Henze et al., 2020; Streit et al., 2014) was completed in the afternoon by *n* = 66 healthy and young (18-33 years) subjects (35 males, 31 females taking oral contraceptives; this sample was first described elsewhere, 11). This exploratory study indicated sex-specific associations between brain structure and increases in cortisol levels, particularly in striatal and frontal regions. With the current study, we aimed at confirming these findings by including additional healthy Scan*STRESS*-samples (Bärtl et al., 2024; Giglberger et al., 2023; Konzok et al., 2021; Speicher et al., 2023), increasing the sample size to *n* = 291 subjects (134 males, 157 females), and further extending them with additional analysis. The present analysis further comprises two parts: a confirmatory analysis, where we reapplied PALM to validate the initial findings (Henze et al., 2023) on a much larger sample, and an exploratory analysis, where we employed a data-driven, hypothesis-free whole-brain approach using Freesurfer’s vertex-wise analysis and machine learning (ML) to investigate the link between cortisol and brain structure on a large scale.

## METHODS AND MATERIALS

### Subjects

This study is based on samples of five independent studies that implemented the Scan*STRESS* protocol on the same scanner with identical parameters, and used the same assay for cortisol determination across all studies (Bärtl et al., 2024; Giglberger et al., 2023; Henze et al., 2020; Konzok et al., 2021; Speicher et al., 2023). From the Regensburg Burnout Project sample, only healthy subjects were included (Bärtl et al., 2024). Additionally, we included data exclusively from subjects who underwent the original version of Scan*STRESS* (Speicher et al., 2023). In total, *n* = 291 subjects (18 − 62 𝑦𝑒𝑎𝑟𝑠, 𝑚𝑒𝑎*n* 𝑎𝑔𝑒 26.16 *±* 8.95) were analyzed, comprising 134 males (18 − 60 𝑦𝑒𝑎𝑟𝑠, 𝑚𝑒𝑎*n* 𝑎𝑔𝑒 26.65 *±* 8.23) and 157 females (18 − 62 𝑦𝑒𝑎𝑟𝑠, 𝑚𝑒𝑎*n* 𝑎𝑔𝑒 25.74 *±* 9.53). Of those female subjects, *n* = 81 were tested during their luteal phase of the menstrual cycle, *n* = 64 were taking hormonal contraceptives, and *n* = 12 were post-menopausal.

Subjects were recruited via flyers and social media platforms. General exclusion criteria across all studies comprised: self-reported history of one or more current psychiatric, neurological, or endocrine disorders; treatment with psychotropic drugs or other medications affecting central nervous system or endocrine function; daily alcohol consumption; incompatibilities with MRI (e.g., metal parts, pregnancy); regular night shifts. Study-specific exclusion criteria are detailed in the original research reports. All subjects provided written informed consent and received financial compensation for their participation. All studies were approved by the local ethics committee of the University of Regensburg.

### General procedure

The test protocol sequence, an overview of ScanSTRESS, and its block design are shown in Figure 1. Test sessions took place between 1 and 6 pm to minimize the influence of the circadian rhythm of cortisol secretion (Kudielka et al., 2004; Zänkert et al., 2019). Subjects arrived at the laboratory 75 minutes before the stress induction followed by a relaxation phase after the first saliva sample was taken to measure cortisol. A total of ten saliva samples were collected (−75, −15, −1, +15, +30, +50, +65, +80, +95, 𝑎*n*𝑑 + 110 𝑚𝑖*n*) using "cortisol salivettes" (Sarstedt, Nuembrecht, Germany); three before, one during, and further six after Scan*STRESS*, with sample -1 referring to stress onset. The saliva samples were analyzed using a time-resolved fluorescence immunoassay with fluorometric endpoint detection (dissociation-enhanced lanthanide fluorescence immunoassay, DELFIA, 46) with an intra-assay coefficient of variation between 4.0 − 6.7% and an inter-assay coefficient of variation between 7.1 − 9.0%.

**Figure 1.**
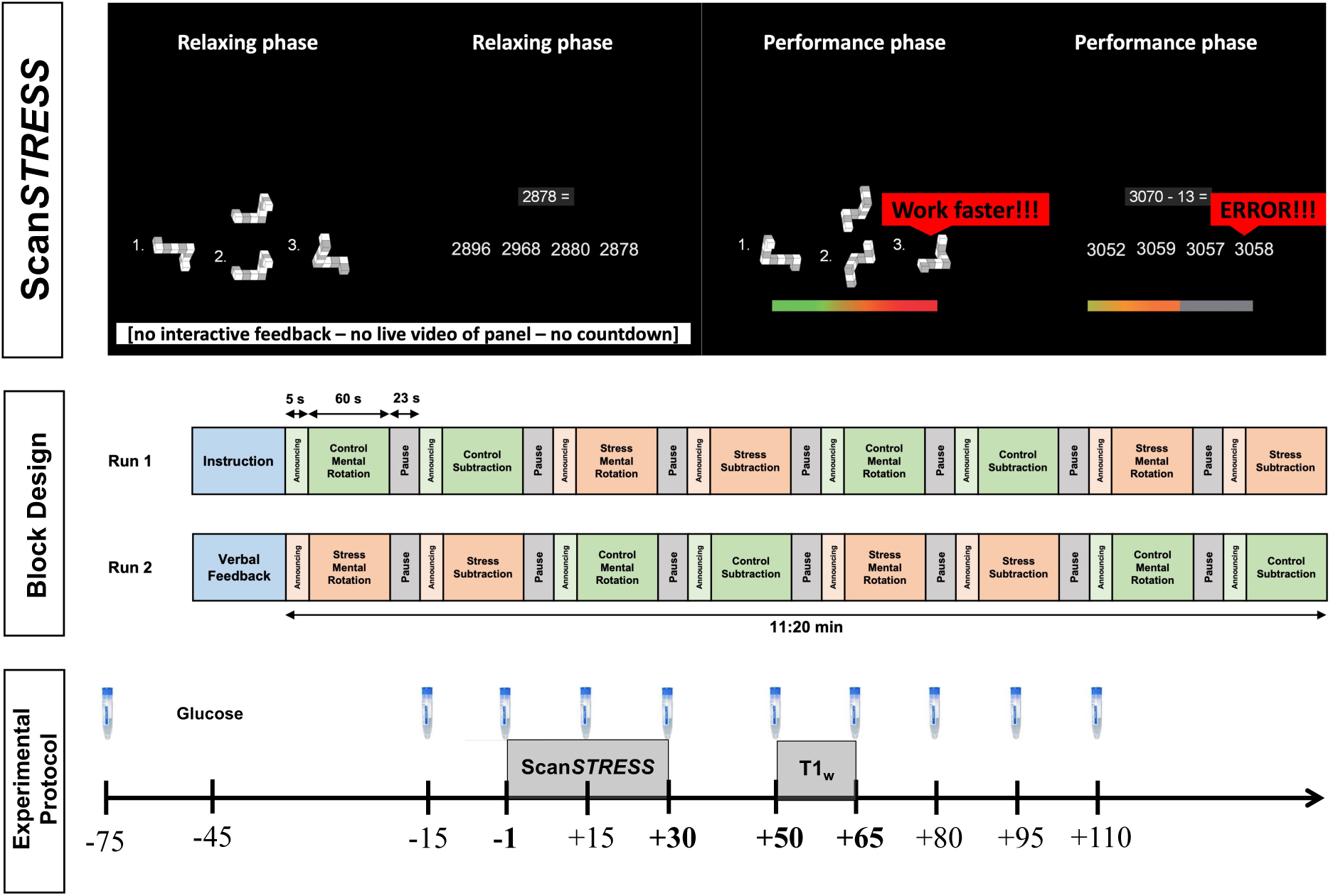
Experimental protocol. Illustration of ScanSTRESS and its block design, repeated collection of salivary cortisol, glucose administration, and acquisition of ScanSTRESS- responses as well as T1-weighted (T1w-) images.

The Scan*STRESS* paradigm is a block-designed protocol with alternating stress and control blocks presented in two runs. During control blocks, subjects perform simple figure and number matching tasks, while stress blocks involve rotation and arithmetic tasks under the supervision of a panel in professional attire, under time pressure, and with constant negative feedback. We used an improved version of the original protocol (Streit et al., 2014) across all included studies to enhance the stress-induced responses without changing the paradigm (Henze et al., 2020). This improvement included: First, a 45-minute relaxation phase was introduced to ensure low baseline cortisol levels, along with a detailed explanation of the scanning procedure to reduce pre-scan anxiety (McGlynn et al., 2007; Thorpe et al., 2008). Second, subjects received a sugary drink (75g glucose in 200ml herbal tea) to boost cortisol reactivity (Zänkert et al., 2020). Third, the transition from relaxation to stress was shortened to under 10 minutes. After Scan*STRESS* exposure, study-specific acquisitions were conducted (i.e., a resting state sequence in all studies except one in which a moral decision paradigm was conducted, 43), followed by an anatomical assessment for all subjects.

### Data analysis

To determine potential relationships between acute cortisol stress responses and neural structural measures, we preprocessed both variables as follows. A cortisol increase was defined as the difference between the peak cortisol level (𝑠𝑎𝑚𝑝𝑙𝑒 + 30, +50, +65) and the pre-stress cortisol level (𝑠𝑎𝑚𝑝𝑙𝑒 − 1), as marked in bold in Figure 1. We focused on cortisol increase as it captures acute stress exposure and HPA axis reactivity-latency (Dickerson & Kemeny, 2004), while area under the curve measures includes pre-stress and recovery phases, reflecting total cortisol release.

Neuroimaging data were collected using a Siemens MAGNETOM Prisma 3𝑇 MRI (Siemens Healthcare, Erlangen, Germany) with a 64-channel head coil. Two series of planar BOLD echo sequences (Scan*STRESS*), multiband resting state (results to be published), and anatomical sequences were performed. Anatomical measurements included a 3D structural T1_w_- image (𝑇1/𝑇𝑅/𝑇𝐸 = 1200/2400/2.18𝑚𝑠, 𝑓𝑙𝑖𝑝 𝑎*n*𝑔𝑙𝑒 = 8, 𝑑𝑖𝑠𝑡𝑎*n*𝑐𝑒 𝑓𝑎𝑐𝑡𝑜𝑟 = 50%).

Anatomical images were initially segmented using SynthSeg, a convolutional neural network-based technique robust to variations in contrast and resolution (Billot et al., 2023). Further preprocessing was conducted using ENIGMA HALFpipe (Waller et al., 2022), which integrates fMRIprep, FSL, and FreeSurfer. FreeSurfer was used for cortical parcellation and subcortical segmentation of the T1_w_-volume, including estimation of TBV (“BrainSegVolNotVent”). Preprocessing steps included non-brain tissue removal, automatic Talairach transformation, segmentation, intensity normalization, gray/white matter boundary tessellation, topology correction, and surface shaping. Additionally, the structural output of each subject was visually inspected before extracting subcortical volume, cortical thickness, and surface area measurements of pre-limbic structures for each hemisphere using asegstats2table and aparcstats2table.

In confirmatory analysis, PALM (Winkler et al., 2014) was used to investigate potential relationships between individual cortisol stress responses and structural measures of the limbic system. As described in our previous study (Henze et al., 2023), the following ROIs were selected based on the current literature on the interaction between neural substrates and acute HPA axis responses (Harrewijn et al., 2020; Van Oort et al., 2017): volumes were extracted for the thalamus, striatum (ncl. caudatus, ncl. accumbens, and putamen), hippocampus, and amygdala. Cortical thickness and surface area were measured for the cingulate cortex (CC: rostral anterior cingulate cortex, rACC; caudal anterior cingulate cortex, cACC; and posterior cingulate cortex, PCC), parahippocampus, the orbitofrontal cortex (OFC: lateral OFC, lOFC; and medial OFC, mOFC), insula, and precuneus. Note that, we use the terminology based on Destrieux- and Desikan-Killiany atlases (Desikan et al., 2006; Destrieux et al., 2010). However, in other literature (Shackman et al., 2011), the rACC is often split into subgenual and pACC, the cACC is called anterior mid-CC (aMCC) or commonly dACC, and the PCC is referred to as posterior mid-CC (pMCC). Six models were tested, where lateralized structural measures (volume, thickness, and surface area) of the left and right hemispheres were separately used as dependent variables, given the assumption that HPA axis regulation may be lateralized (Cerqueira et al., 2008) and following our preliminary results (Henze et al., 2023). Since sex, age, and TBV can influence structural brain measures (Giedd et al., 1999; Moog et al., 2021), these variables were included as covariates, with TBV considered only in volumetric models. In addition, menstrual cycle phase, hormonal contraceptive use, and reproductive status also influence brain structure (Brønnick et al., 2020; B. Pletzer et al., 2023; Song et al., 2023). Therefore, each of the six models with *lateralized structural measures* as dependent variables included *cycle* (male, luteal phase, contraception, post-menopause; dummy-coded with male as reference), *age* (continuous), and *TBV* (continuous, where appropriate) as control variables, along with individual *(sex-specific) cortisol increases* (continuous) as predictors. This grouping variable was implemented as follows, for instance, male cortisol increase was coded to contain the actual cortisol increase value for male subjects and 0 for female subjects; the same procedure was applied for female cortisol increase, but in reverse order. This approach allowed us to correct for and investigate the possible influence of sex-specific cortisol responses. To test for the general relationship between cortisol increase and brain structure (independent of sex), the influence of both variables, male and female cortisol increase, was tested simultaneously (see Supplementary Methods for tested statistical model, 14). We did not create a variable to differentiate cortisol increases based on the *cycle* categories, even though such differences were reported (Zänkert et al., 2019), because although there was significant variation among groups 𝐹(3, 287) = 13.70, 𝑝 < .000), this was driven by significant differences between males compared to two female categories (luteal phase, 𝑝 < .000; contraceptive users, 𝑝 < .000), but not among the female categories (𝑝 > .637) (Supplementary Table S1). Multiple comparisons were corrected using the False Discovery Rate (FDR) -fdr option.

In the first stage of exploratory analyses, FreeSurfer’s (version 7.4.1) vertex-wise general linear model (GLM)-based group analyses were performed to investigate the relationship between sex-specific cortisol responses and structural measures of the entire cortex without prior assumptions about regions. Subjects’ cortical surface was first mapped onto the fsaverage template, smoothed with a 10𝑚𝑚 full width at half-maximum kernel, and concatenated into a single dataset for each hemisphere (mris_preproc). In the GLM (mri_glmfit, Different Offset, Different Slope), *volume*, *thickness*, and *surface area* were dependent variables, while *sex-specific cortisol increase* was the predictor, and the confounding effects of *cycle* and *age* were controlled. As in PALM-analyses, sex-independent and sex-specific cortisol increase were evaluated separately using corresponding contrast matrices in vertex-based analyses. Monte Carlo Simulation cluster analyses at a significance level of 𝑝 = .050 were used to correct for multiple comparisons (mri_glmfit-sim –2spaces). The cluster-forming threshold was set to 3.0 to prevent inflated false positives (Greve & Fischl, 2018).

As further exploratory analyses, an ML approach, ridge regression, was employed to predict subjects’ cortisol stress response from their volume, thickness, and surface area measures (158 features in total). Although the GLM is a standard method for localizing the relationship between cortisol increase and structural brain measures, evaluating the generalizability of these associations is not straightforward. However, ML inherently enables out-of-sample evaluation, which is essential for evaluating generalizability (Kriegeskorte et al., 2009). Additionally, as the generative process is reversed in ML compared to GLM, it allows for the extraction of complex multivariate patterns in brain measures to predict cortisol response, thereby enabling the identification of the physiological stress state at the subject level. Briefly, ridge regression was fit and evaluated within a ten-repeated nested cross-validation framework: Ten outer folds for model evaluation and five inner folds for hyperparameter optimization. Ridge regression is a type of multivariate linear regression that incorporates regularization, which shrinks the coefficients of uninformative features, thereby making it more robust to multicollinearity (Hoerl & Kennard, 1970). Cross-validation is a standard approach to evaluating the predictive performance of a model by splitting the data into training and test sets recursively. As cross-validation is a stochastic approach (i.e., the data is split randomly), we repeated the cross-validation 10 times to reduce arbitrariness (Varoquaux et al., 2017). Additionally, the potential confounding effect of *cycle* was corrected using a cross-validated confound-correction technique to avoid any possible data leakage (Snoek et al., 2019). *Age* was not corrected as it was not found to be associated with *cortisol increase* (𝐹(289, 1) = .819, 𝑝 = .366). The model performance was evaluated using correlation coefficient (*r*) and tested against a null distribution using permutation testing (Ojala & Garriga, 2009) with 1000 permutations for significance. Shapley additive explanations (SHAP) were used to evaluate the contributions of brain measures to the prediction performance (Lundberg & Lee, 2017). Details of model interpretation are given in Supplementary Methods.

## RESULTS

Sample characteristics of sex-specific cortisol increase and TBV are given in Supplementary Table S2. Female subjects showed a significantly lower cortisol response to the stress stimulus than males (𝑡(290) = −6.291, 𝑝 = .001) and exhibited lower TBV (𝑡(290) = −14.226, 𝑝 =.001).

### Permutation Analysis of Linear Models (PALM) results

The significant results from PALM-analyses are presented in Table 2, whereas full results are provided in Supplementary Tables S4-9.

**Table 2.**
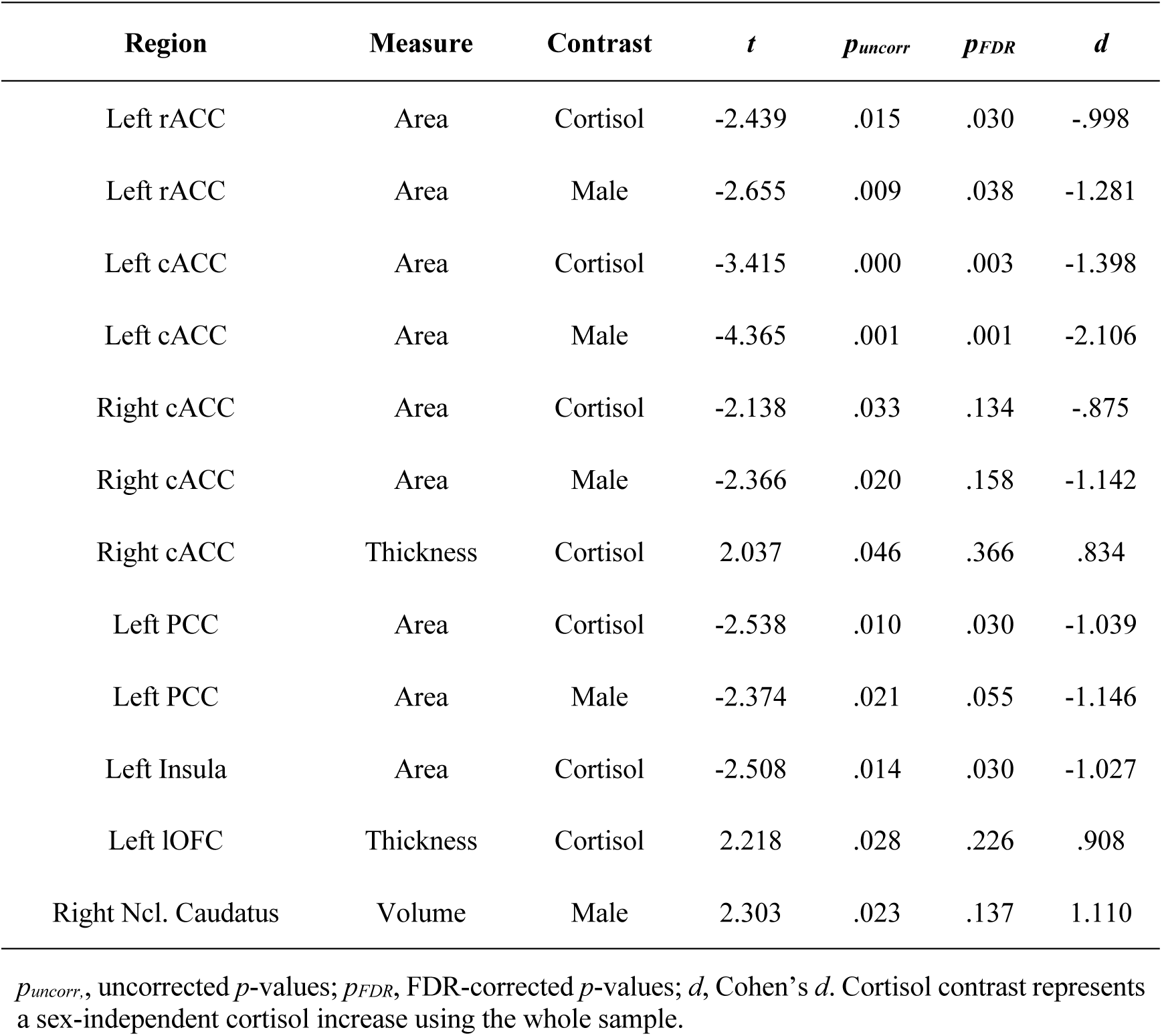
Summary of significant PALM Calculations for surface area, thickness, and volume.

The CC was found to be consistently negatively associated with *cortisol increase* across hemispheres and certain brain measures. Specifically, a reduced *surface area* in the left hemisphere of rACC, cACC, PCC, and insula was significantly associated with sex- independent *cortisol increase*. Of these regions, *surface area* of rACC, cACC, and PCC was also negatively associated with *cortisol increase* in males. In the right hemisphere, only the *surface area* of cACC showed a marginally significant association with sex-independent and male-specific *cortisol increase*, though, these did not survive FDR-correction. For *cortical thickness*, positive relationships between sex-independent *cortisol increase* and right cACC as well as left lOFC were observed. In the subcortex, a marginally positive association between right ncl. caudatus *volume* and *cortisol increase* in males was found. However, none of these associations withstand FDR-correction. Figure 2 depicts scatterplots of associations between (sex-specific) cortisol increases and structural brain measures.

**Figure 2.**
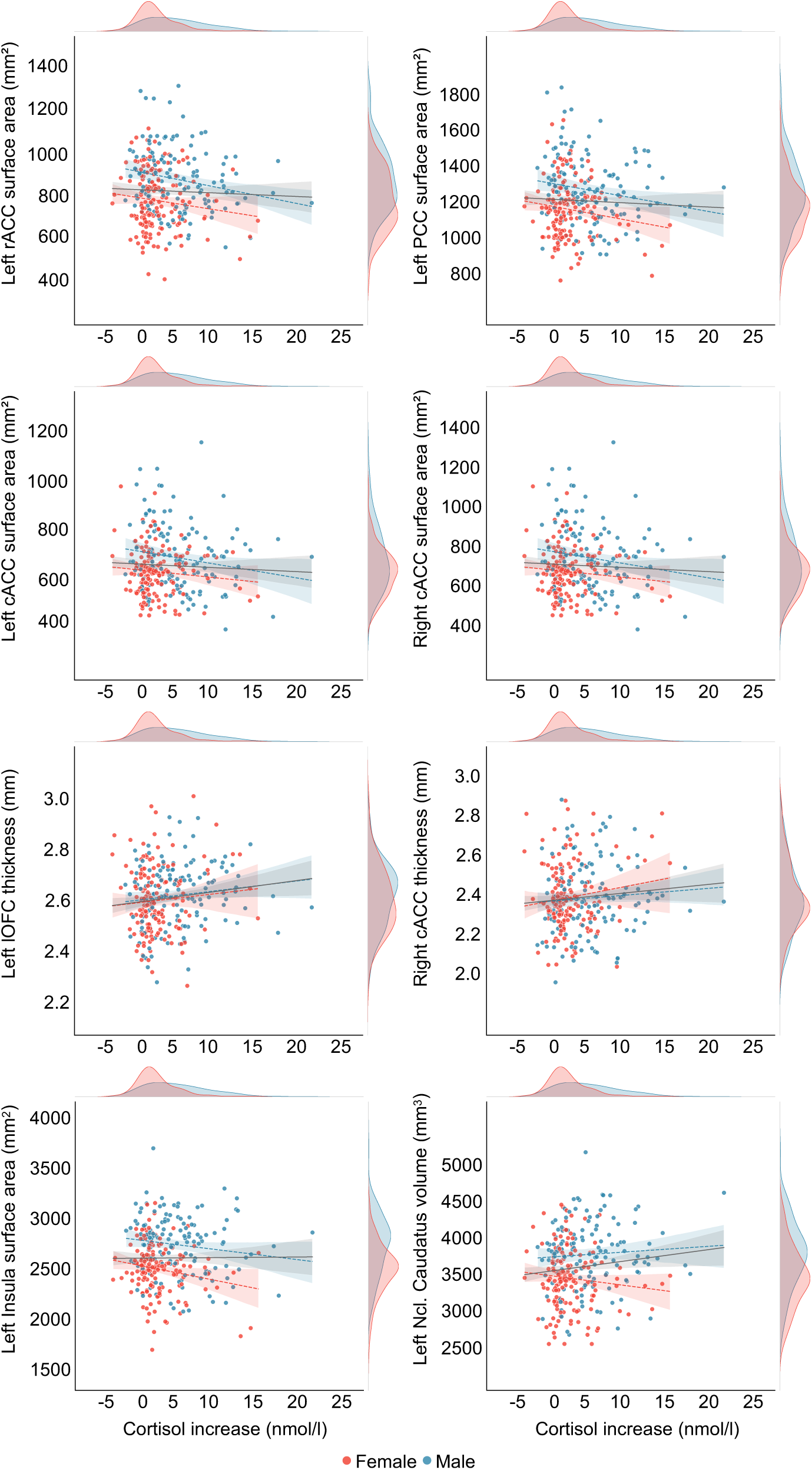
Scatterplots of associations between individual cortisol increase (nmol/L) and structural brain measures of the rostral anterior cingulate cortex (rACC), posterior cingulate cortex (PCC), caudal anterior cingulate cortex (cACC), lateral orbitofrontal cortex (lOFC), insula, and ncl. caudatus. Male and female subjects are represented by blue and red colors, respectively. The linear gray line represents a model fit to the total sample, while sex-specific models are depicted with respective colored dotted lines. The density plots illustrate the distribution of cortisol increase and the brain measures for each sex.

### FreeSurfer’s Vertex-Wise Group Analyses

In line with the results from PALM-analysis, vertex-wise group analyses revealed a significant negative association between *surface area* of the left cACC and *cortisol increase* in males, which was also observed in volume (Figure 3 and Supplementary Table 3).

**Figure 3.**
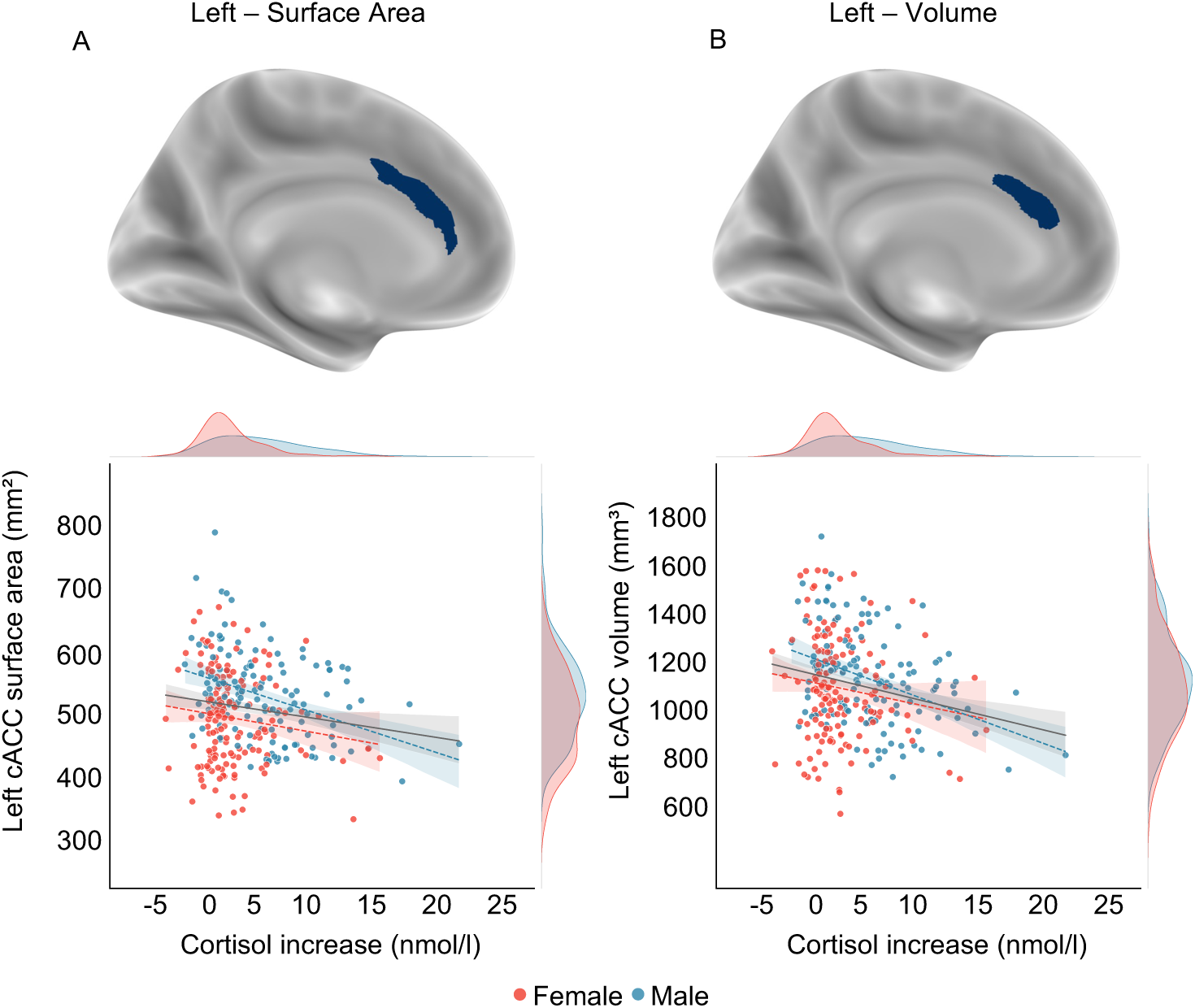
Whole-brain vertex-wise association results surviving multiple comparisons correction (cluster-wise *p_corrected_* < .05) and scatter plots depicting corresponding associations with cortisol increase, which are only negative. In the scatter plots, sex groups are depicted with distinct colors. The linear gray slope represents a model fit to the total sample, while sex-specific models are depicted with colored dotted lines. The density plots represent the distribution of cortisol increase and the brain measures for each sex. A lower surface area and volume of the caudal anterior cingulate cortex (cACC) were significantly associated with a greater cortisol increase in males.

### Machine-Learning Analysis

Ridge regression with hyperparameter optimization significantly predicted subjects’ *cortisol increase* based on their cortical *surface area* and *thickness*, as well as subcortical *volume* measures, with out-of-sample scores yielding a correlation coefficient of 𝑟 = .124 (𝑝 = .039).

Figure 4 depicts the most informative structural measures for predicting individuals’ *cortisol increase*. Most of the brain measures reported in the previous analyses were also found to be informative in predicting *cortisol increase*, such as cACC, PCC, lOFC, as well as ncl. caudatus, along with additional areas from frontal, temporal, and limbic regions. Left cACC *surface area* was found to be the most informative feature in predicting *cortisol increase*. Additionally, the direction of the impact of brain measures on outcome cortisol increase values was found to be mostly negative, indicating a negative association between these measures and cortisol increase. Moreover, these informative brain measures were found to be stable across cross-validation folds, indicating their generalizability (Supplementary Figure S1).

**Figure 4.**
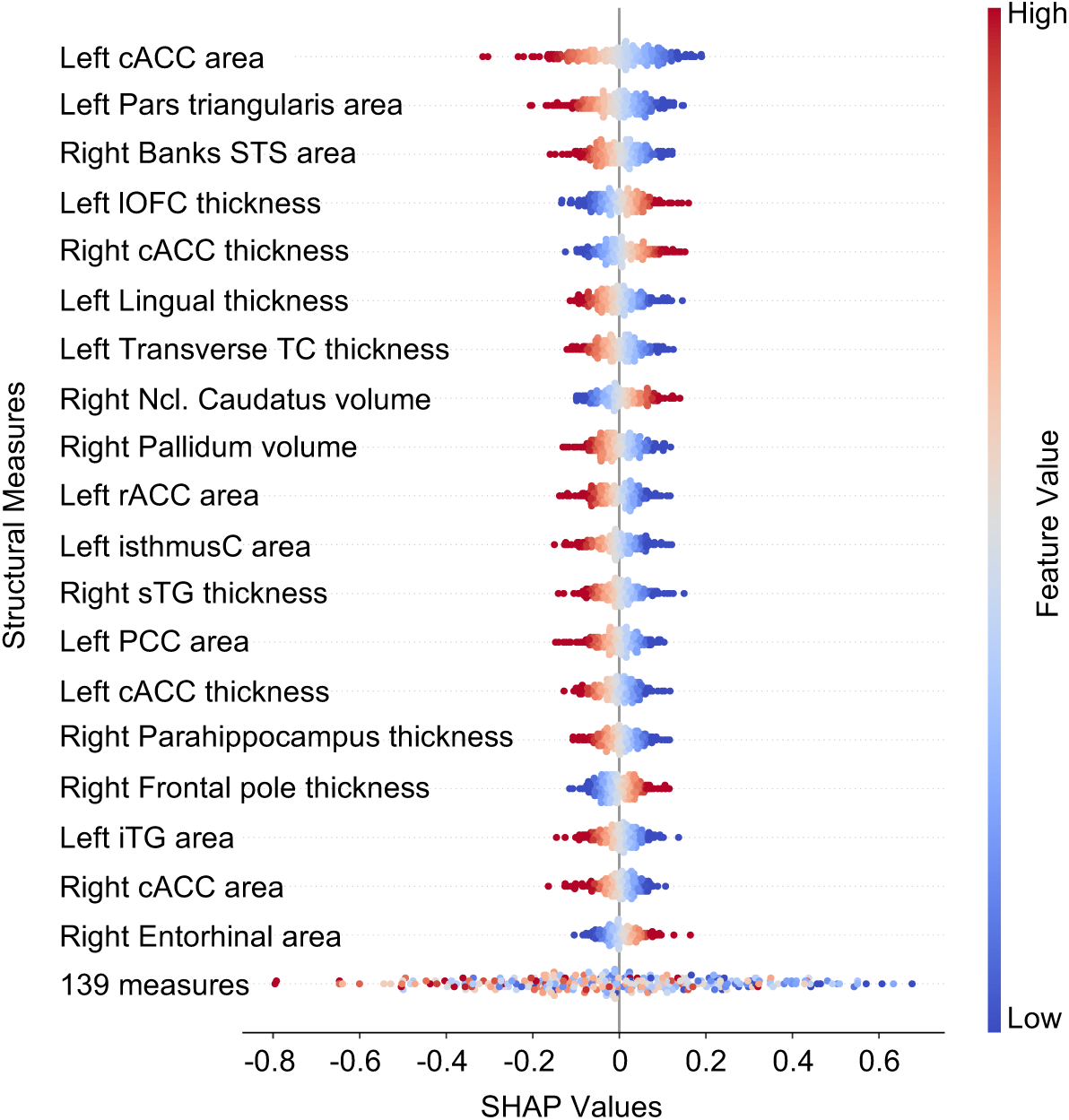
The most informative structural measures, as measured by SHAP values derived from a trained ridge regression model, are indicative of their impact on the final prediction scores. The direction and extent of the SHAP values indicate the direction and magnitude of the features’ effects on the final prediction scores. The feature values are represented by colors, with blue indicating low values and red indicating high values. cACC, caudal anterior cingulate cortex; STS, superior temporal sulcus; lOFC, lateral orbitofrontal cortex; TC, temporal cortex; Ncl. Caudatus, nucleus caudatus; isthmusC, isthmus cingulate; sTG, superior temporal gyrus; PCC, posterior cingulate cortex; iTG, inferior temporal gyrus; 139 regions, the sum of remaining 139 brain measures.

## DISCUSSION

The present study revealed consistent negative associations between structural measures of the CC and acute cortisol increases in response to psychosocial stress, across three independent analyses: ROI-based PALM-analysis, whole-brain with Freesurfer’s vertex-wise group analysis, and out-of-sample prediction using ridge regression (i.e., regularized multivariate linear regression). Specifically, smaller *surface area* of the left rACC, cACC, and PCC, as well as the right cACC, was significantly associated with higher *cortisol increase*, regardless of *sex*. These effects were also found in the male sub-sample. The negative relationship between left cACC *surface area* and *cortisol increase* in males was further validated through whole-brain analysis and extended to the volume measure of the cACC. Notably, the *surface area* of left cACC also emerged as the most promising predictor of stress-induced cortisol increases in prediction analysis (independent of *sex* and *cycle*). These findings highlight the predominantly negative impact of cingulate surface area on stress-induced cortisol release. Additionally, our results suggest that structural measures of other cortical and subcortical areas (e.g., lOFC, insula, ncl. caudatus) are also associated with or predictive of acute cortisol increases in response to stress.

These findings complement previous task-based and rsFC-studies showing that the CC is closely linked to the HPA axis and stress processing (Berretz et al., 2021; Harrewijn et al., 2020; Noack et al., 2019; Qiu et al., 2022; Van Oort et al., 2017): For instance, Scan*STRESS* studies have reported positive associations between cortisol responses and activations in the PCC (Henze et al., 2020) and cACC (65; referred to as dACC), as well as in the pACC (66, acting as rACC-cACC transition zone). Additionally, a study using another psychosocial stress paradigm – the Montreal Imaging Stress Task (MIST, (Dedovic et al., 2005; J. C. Pruessner et al., 2008) – confirmed this positive relationship for the pACC (Boehringer et al., 2015). Another study showed that activation-habituation in the bilateral ACC was accompanied by blunted cortisol release induced by Scan*STRESS* and MIST (Y. Liu et al., 2023). Interestingly, Neurexan® (Nx4), a natural pharmacological agent, has been shown to reduce cortisol responses to psychosocial stress (Doering et al., 2016) and lower stress-induced activation, particularly in the supracallosal ACC (functionally part of the cACC, 70). This study also found a significant association between trait anxiety and the Nx4-effect on stress-induced rsFC- changes. Notably, subjects with above-average trait anxiety exhibited marked improvement in stress-related rsFC-changes between the right amygdala and pACC.

The relevance of cingulate structures is also reflected in the few studies on structural measures and HPA axis reactivity: For example, rapid volumetric changes following acute psychosocial stress induction (TSST) have been observed in anterior and mid-cingulate regions with increases in volume being positively related to stress-elicited state anxiety elevations (Uhlig et al., 2023). Moreover, reduced volume and heightened stress-related brain activity in the pACC were linked to a lower cortisol awakening response (CAR). Interestingly, rsFC between the pACC and hypothalamus – the primary regulator of HPA axis activity – also exhibited a negative association with the CAR (Boehringer et al., 2015).

Taken together, both functional and structural studies suggest that the CC plays a crucial role in emotional and cognitive appraisal of stressful stimuli. Positive correlations between stress-induced cortisol responses and the activation of CC-structures, as well as their functional connections to other stress-relevant brain regions, emphasize its importance as an integrator of stress processes. Specifically, our findings suggest that structural differences, particularly smaller surface area or volume (Boehringer et al., 2015), may impair the ability to effectively regulate stress, as reflected in increased cortisol responses to acute stress. This positions the CC as a potential biomarker for stress susceptibility and HPA axis reactivity, which is relevant for stress-related disorders (Hinojosa et al., 2019; Huang et al., 2024; Misquitta et al., 2021; Simmons et al., 2008; Yang et al., 2022; Yucel et al., 2008; H. Zhang et al., 2024). Furthermore, initial studies suggest that markers for resilience and treatment responses can be localized in these neural regions (Alagapan et al., 2023; Carnevali et al., 2018; Pizzagalli et al., 2001; Webb et al., 2018). However, it should be noted that different patterns of effects and functions may prevail depending on the specific subregion, as confirmed by inconsistent anatomical labelling in studies (Shackman et al., 2011). This highlights the need for specific investigations of the stress-specific functionality of cingulate substructures (Bubb et al., 2018; Rolls, 2019).

Moreover, the current results support and extend our previous findings (Henze et al., 2023): In this larger sample, we also observed a positive association between frontal cortex morphology and stress-induced cortisol release, specifically for the thickness of right cACC and left lOFC. This suggests that frontal regions may play a key role in stress perception, engaging in top-down processing that incorporates limbic circuits (e.g., amygdala, hippocampus) to regulate stress response systems like the HPA axis (Herman et al., 2005, 2016; Hermans et al., 2024; Jankord & Herman, 2008; Van Oort et al., 2017). We once again observed a positive association between right ncl. caudatus volume and cortisol levels in males, further reinforcing the existence of sex-specific patterns in hormonal stress responses involving striatal structures (Henze et al., 2021). The difference between male and female brain-stress correlations was, however, not significant, likely due to the broader sample composition and the need for statistical control of the variable *cycle*. It is well-established that both the cortisol stress response and brain morphology are influenced by factors such as menstrual cycle phase, hormonal contraception, and reproductive status (Brønnick et al., 2020; Childs et al., 2010; Goldstein et al., 2010; Hidalgo-Lopez et al., 2020; Klusmann et al., 2022; B. Pletzer et al., 2010, 2018; B. Pletzer, Harris, & Hidalgo-Lopez, 2019; B. Pletzer, Harris, Scheuringer, et al., 2019; B. A. Pletzer & Kerschbaum, 2014; Rincon-Cortes et al., 2019; Yoest et al., 2018). For the structural data, we therefore applied a non-binary correction (i.e. *cycle*), but we distinguished cortisol responses dichotomously between males and females, as there were no significant differences between female subgroups regarding cortisol, despite potential underlying variations (Klusmann et al., 2022; Zänkert et al., 2019). This emphasizes the need for larger studies on female subgroups and more comprehensive data in this area.

Our ML-model achieved modest but significant prediction performance for cortisol increases based on structural measures. While this is the first ML-study in this context, similar low correlations have been reported in studies predicting PTSD-symptoms and cortisol from rsFC and acute stress responses (W. Zhang et al., 2020, 2022). Despite these parallels, our model’s 𝑟 is not high enough for clinical application and should be interpreted cautiously. Additionally, SHAP-analysis highlighted the importance of surface area alongside volume and thickness, consistent with the findings from PALM and whole-brain analyses. Re-fitting the model without surface area resulted in significantly lower prediction performance, emphasizing its necessity in future stress studies (Supplementary Table S10). This is further supported by the observed negative association between insular *surface area* and *cortisol increase*. Although the insula has shown high relevance in functional stress studies (Berretz et al., 2021; Noack et al., 2019; Van Oort et al., 2017), it has so far been neglected in structural studies. Our findings indicate that its involvement may correspond to that of the CC (Bahamonde et al., 2023; Hu et al., 2018; Kang et al., 2017; Shao et al., 2018).

In conclusion, our findings demonstrate that surface area, thickness, and volume of both cortical and subcortical structures can serve as predictors of hormonal stress responses, expanding our understanding of key brain regions and the importance of various structural metrics. Future studies should leverage even larger samples to capture more variance in sex-and potentially also gender-specific effects and facilitate the application of advanced ML- methods. Considering these findings, smaller surface areas of the CC may favor a higher HPA axis stress reactivity. Specifically, the left cACC emerges as a crucial predictor of stress- induced cortisol release, highlighting its potential as a biomarker for stress-related diseases.

## DATA AVAILABILTY STATEMENT

The datasets generated during and/or analyzed during the current study are available from the corresponding author on reasonable request. Analysis codes can be found at https://github.com/eminSerin/stress-fs-paper. A repository of studies that have already used and published this data is available here: https://osf.io/echja/.

## AUTHOR CONTRIBUTIONS

**Emin Serin**: Conceptualization, Methodology, Software, Validation, Formal analysis, Data Curation, Writing – Original Draft, Writing – Review & Editing, Visualization. **Lea Sophie Schill**: Data Curation, Writing – Review & Editing. **Christoph Bärtl**: Investigation, Writing – Review & Editing. **Marina Giglberger**: Investigation, Writing – Review & Editing. **Julian Konzok**: Investigation, Writing – Review & Editing. **Hannah L. Peter**: Investigation, Writing – Review & Editing. **Nina Speicher**: Investigation, Writing – Review & Editing. **Ludwig Kreuzpointner**: Writing – Review & Editing. **Brigitte M. Kudielka**: Writing – Review & Editing. **Stefan Wüst**: Writing – Review & Editing. **Henrik Walter**: Resources, Writing – Review & Editing, Supervision, Funding acquisition. **Gina-Isabelle Henze**: Conceptualization, Methodology, Software, Validation, Formal analysis, Investigation, Resources, Data Curation, Writing – Original Draft, Writing – Review & Editing, Visualization, Project administration, Funding acquisition.

## ORCID

Emin Serin: https://orcid.org/0000-0002-3570-3027

Christoph Bärtl: https://orcid.org/0000-0002-0349-6254

Marina Giglberger: https://orcid.org/0000-0002-1828-8474

Julian Konzok: https://orcid.org/0000-0002-4232-4105

Nina Speicher: https://orcid.org/0000-0003-2725-6498

Ludwig Kreuzpointner: https://orcid.org/0000-0003-1391-0807

Brigitte M. Kudielka: https://orcid.org/0000-0002-8349-926X

Stefan Wüst: https://orcid.org/0000-0002-2315-8949

Henrik Walter: https://orcid.org/0000-0002-9403-6121

Gina-Isabelle Henze: https://orcid.org/0000-0002-3397-1351

