## Supplementary Material for "Linking brain structure to stress reactivity: Cingulate surface area predicts acute cortisol responses"

### Supplementary Methods

**Statistical model tested with PALM:**

| Contrast | Male cortisol | Female cortisol | Cycle | Age | TBV* |
|---|---|---|---|---|---|
| Cortisol | 1 | 1 | 0 | 0 | 0 |
| Male | 1 | 0 | 0 | 0 | 0 |
| Female | 0 | 1 | 0 | 0 | 0 |
| Male > Female | 1 | -1 | 0 | 0 | 0 |
| Female > Male | -1 | 1 | 0 | 0 | 0 |

*Total Brain Volume (TBV) was only included in volumetric models.

**Feature Importance Analysis:** Shapley additive explanations (SHAP) were utilized to identify contributing brain measures that drive cortisol increase prediction (Lundberg & Lee, 2017). SHAP is a model-agnostic method that measures the marginal contribution of each feature to a model prediction, providing a straightforward way to rank and visualize features based on their contribution. As ridge regression was employed, the linear SHAP estimator was used. We calculated SHAP values in two ways. First, after model training and evaluation, we fit the model using the entire sample and estimated SHAP values. This provides a single set of straightforwardly interpretable SHAP values (Figure 3). Second, we estimated SHAP values on the test set of each outer cross-validation fold, resulting SHAP values with a size of S x F in each fold, where S represents subjects within test fold and F represents brain measures. Within each fold, we took median absolute SHAP value for each brain measure, providing a final matrix of CV folds x brain measures. This allows us to evaluate the generalizability and variability of brain measure importance (Supplementary Figure 1). We identified important features based on the mean absolute SHAP value in each analysis and reported the top 20 informative brain measures.

### Supplementary Tables

**Table S1.** Post-hoc comparisons of cortisol increase among sex and cycle groups.

| Groups | | Mean Difference | *p* |
|---|---|---|---|
| Males | Luteal Phase | -2.891 | .001 |
| Males | Contraceptive | -2.206 | .001 |
| Males | Post-Menopause | -2.150 | .167 |
| Luteal Phase | Contraceptive | .685 | .637 |
| Luteal Phase | Post-Menopause | .741 | .900 |
| Contraceptive | Post-Menopause | .056 | 1.0 |

One-Way ANOVA: $F(3, 287) = 13.70, p < .001$). Post-hoc comparisons were done using Tukey's HSD.

**Table S2.** Sample characteristics of cortisol increase and total brain volume by sex.

| | Mean | *SD* | *t* | *p* |
|---|---|---|---|---|
| Cortisol increase (nmol/L) | | | | |
| Female | 1.692 | 2.777 | -6.291 | .001 |
| Male | 4.248 | 4.108 | | |
| Total brain volume ($mm^3$) | | | | |
| Female | 1084907.554 | 77667.106 | -14.226 | .001 |
| Male | 1219128.567 | 83121.059 | | |

**Table S3.** Significant clusters based on vertex-wise analysis of cortisol increase x structural brain measures effect.

| Region | Measure | Contrast | Max. Vert. Sig. -log(*p*) | *p* | Cluster Size ($mm^2$) | MNI Coordinates (x, y, z) |
|---|---|---|---|---|---|---|
| Left cACC | Volume | Male | -4.855 | .004 | 326.73 | -8.2, 31.1, 20.3 |
| Left cACC | Area | Male | -4.522 | .007 | 458.99 | -5.9, 24.3, 18.0 |

Max. Vert. Sig., maximum vertex significance; MNI, Montreal Neurological Institute

**Table S4.** Results of PALM calculations for cortical surface area of the left hemisphere.

| Contrast | rACC | cACC | PCC | Parahippocampus | lOFC | mOFC | Insula | Precuneus |
|---|---|---|---|---|---|---|---|---|
| **Cortisol** | | | | | | | | |
| $T$ | **-2.439** | **-3.415** | **-2.538** | -1.772 | -1.110 | -.282 | **-2.508** | -1.097 |
| $p_{uncorr}$ | **.015** | **.000** | **.010** | .075 | .263 | .783 | **.014** | .273 |
| $p_{FDR}$ | **.030** | **.003** | **.030** | .120 | .312 | .783 | **.030** | .312 |
| $d$ | **-.998** | **-1.398** | **-1.039** | -.726 | -.454 | -.116 | **-1.027** | -.449 |
| **Male** | | | | | | | | |
| $t$ | **-2.655** | **-4.365** | **-2.374** | -1.020 | -.601 | -.043 | -1.890 | -.745 |
| $p_{uncorr}$ | **.009** | **.000** | **.021** | .316 | .555 | .967 | .060 | .458 |
| $p_{FDR}$ | **.038** | **.001** | .055 | .505 | .634 | .967 | .120 | .611 |
| $d$ | **-1.281** | **-2.106** | -1.146 | -.492 | -.290 | -.021 | -.912 | -.359 |
| **Female** | | | | | | | | |
| $t$ | -1.084 | -1.051 | -1.409 | -1.445 | -.933 | -.317 | -1.723 | -.812 |
| $p_{uncorr}$ | .284 | .290 | .157 | .151 | .356 | .752 | .084 | .414 |
| $p_{FDR}$ | .465 | .465 | .418 | .418 | .473 | .752 | .418 | .473 |
| $d$ | -.543 | -.526 | -.705 | -.723 | -.467 | -.159 | -.862 | -.407 |
| **Male > Female** | | | | | | | | |
| $t$ | -.675 | -1.701 | -.248 | .573 | .403 | .232 | .289 | .222 |
| $p_{uncorr}$ | .507 | .087 | .810 | .569 | .694 | .817 | .769 | .819 |
| $p_{FDR}$ | .819 | .698 | .819 | .819 | .819 | .819 | .819 | .819 |
| $d$ | -.322 | -.812 | -.118 | .274 | .193 | .111 | .138 | .106 |
| **Female > Male** | | | | | | | | |
| $t$ | .675 | 1.701 | .248 | -.573 | -.403 | -.232 | -.289 | -.222 |
| $p_{uncorr}$ | .507 | .087 | .810 | .569 | .694 | .817 | .769 | .819 |
| $p_{FDR}$ | .819 | .698 | .819 | .819 | .819 | .819 | .819 | .819 |
| $d$ | .322 | .812 | .118 | -.274 | -.193 | -.111 | -.138 | -.106 |

$p_{uncorr}$: uncorrected $p$-values; $p_{FDR}$: FDR-corrected $p$-values; $d$: Cohen's $d$; rACC: rostral anterior cingulate cortex; cACC: caudal ACC; PCC: posterior cingulate cortex; lOFC: lateral orbitofrontal cortex; mOFC: medial OFC. Cortisol contrast represents a sex-independent cortisol increase using the whole sample. Significant results are given in bold.

**Table S5.** Results of PALM calculations for cortical surface area of the right hemisphere.

| Contrast | | rACC | cACC | PCC | Parahippocampus | lOFC | mOFC | Insula | Precuneus |
|---|---|---|---|---|---|---|---|---|---|
| **Cortisol** | | | | | | | | | |
| | ***t*** | -1.165 | **-2.138** | -.887 | -.984 | -1.835 | -1.829 | -1.881 | -1.4905 |
| | $p_{uncorr}$ | .254 | **.033** | .368 | .325 | .067 | .067 | .056 | .138 |
| | $p_{FDR}$ | .338 | .134 | .368 | .368 | .134 | .134 | .134 | .221 |
| | ***d*** | -.477 | -.875 | -.363 | -.403 | -.751 | -.749 | -.770 | -.610 |
| **Male** | | | | | | | | | |
| | ***t*** | -1.222 | **-2.366** | -.902 | -.509 | -1.659 | -1.214 | -1.082 | -.609 |
| | $p_{uncorr}$ | .229 | **.020** | .376 | .617 | .104 | .227 | .279 | .541 |
| | $p_{FDR}$ | .446 | .158 | .501 | .617 | .416 | .446 | .446 | .617 |
| | ***d*** | -.590 | -1.142 | -.435 | -.245 | -.800 | -.586 | -.522 | -.294 |
| **Female** | | | | | | | | | |
| | ***t*** | -.552 | -.922 | -.440 | -.844 | -1.061 | -1.375 | -1.533 | -1.395 |
| | $p_{uncorr}$ | .587 | .351 | .651 | .392 | .287 | .171 | .119 | .165 |
| | $p_{FDR}$ | .651 | .522 | .651 | .522 | .522 | .457 | .457 | .457 |
| | ***d*** | -.276 | -.462 | -.220 | -.422 | -.531 | -.688 | -.768 | -.698 |
| **Male > Female** | | | | | | | | | |
| | ***t*** | -.268 | -.637 | -.171 | .385 | -.111 | .403 | .608 | .773 |
| | $p_{uncorr}$ | .796 | .515 | .863 | .701 | .913 | .685 | .542 | .446 |
| | $p_{FDR}$ | .913 | .913 | .913 | .913 | .913 | .913 | .913 | .913 |
| | ***d*** | -.128 | -.304 | -.082 | .184 | -.053 | .192 | .290 | .369 |
| **Female > Male** | | | | | | | | | |
| | ***t*** | .268 | .637 | .171 | -.385 | .111 | -.403 | -.608 | -.773 |
| | $p_{uncorr}$ | .796 | .515 | .863 | .701 | .913 | .685 | .542 | .446 |
| | $p_{FDR}$ | .913 | .913 | .913 | .913 | .913 | .913 | .913 | .913 |
| | ***d*** | .128 | .304 | .082 | -.184 | .053 | -.192 | -.290 | -.369 |

$p_{uncorr}$: uncorrected *p*-values; $p_{FDR}$: FDR-corrected *p*-values; *d*: Cohen's *d*; rACC: rostral anterior cingulate cortex; cACC: caudal ACC; PCC: posterior cingulate cortex; lOFC: lateral orbitofrontal cortex; mOFC: medial OFC. Cortisol contrast represents a sex-independent cortisol increase using the whole sample. Significant results are given in bold.

**Table S6.** Results of PALM calculations for cortical thickness of the left hemisphere.

| Contrast | | rACC | cACC | PCC | Parahippocampus | lOFC | mOFC | Insula | Precuneus |
|---|---|---|---|---|---|---|---|---|---|
| **Cortisol** | | | | | | | | | |
| | ***t*** | -1.073 | -.559 | .035 | -.303 | **2.218** | .738 | -.059 | .318 |
| | $p_{uncorr}$ | .283 | .578 | .974 | .764 | **0.028** | .463 | .952 | .744 |
| | $p_{FDR}$ | .974 | .974 | .974 | .974 | 0.226 | .974 | .974 | .974 |
| | ***d*** | -.439 | -.229 | .014 | -.124 | 0.908 | .302 | -.024 | .130 |
| **Male** | | | | | | | | | |
| | ***t*** | -.108 | -.759 | -.381 | -.548 | 1.287 | .059 | 1.168 | -.135 |
| | $p_{uncorr}$ | .913 | .447 | .704 | .576 | 0.203 | .955 | .242 | .892 |
| | $p_{FDR}$ | .955 | .955 | .955 | .955 | 0.955 | .955 | .955 | .955 |
| | ***d*** | -.052 | -.366 | -.184 | -.264 | 0.621 | .029 | .564 | -.065 |
| **Female** | | | | | | | | | |
| | ***t*** | -1.243 | -.140 | .318 | .023 | 1.801 | .866 | -.917 | .488 |
| | $p_{uncorr}$ | .217 | .887 | .753 | .981 | 0.078 | .390 | .356 | .624 |
| | $p_{FDR}$ | .780 | .981 | .981 | .981 | 0.627 | .780 | .780 | .981 |
| | ***d*** | -.622 | -.070 | .159 | .011 | 0.902 | .433 | -.459 | .245 |
| **Male > Female** | | | | | | | | | |
| | ***t*** | .942 | -.331 | -.480 | -.339 | -0.705 | -.666 | 1.425 | -.438 |
| | $p_{uncorr}$ | .346 | .735 | .633 | .730 | 0.488 | .503 | .152 | .639 |
| | $p_{FDR}$ | .735 | .735 | .735 | .735 | 0.735 | .735 | .735 | .735 |
| | ***d*** | .450 | -.158 | -.229 | -.162 | -0.336 | -.318 | .680 | -.226 |
| **Female > Male** | | | | | | | | | |
| | ***t*** | -.942 | .331 | .480 | .339 | 0.705 | .666 | -1.425 | .438 |
| | $p_{uncorr}$ | .346 | .735 | .633 | .730 | 0.488 | .503 | .152 | .639 |
| | $p_{FDR}$ | .735 | .735 | .735 | .735 | 0.735 | .735 | .735 | .735 |
| | ***d*** | -.450 | .158 | .229 | .162 | 0.336 | .318 | -.680 | .226 |

$p_{uncorr}$: uncorrected *p*-values; $p_{FDR}$: FDR-corrected *p*-values; *d*: Cohen's *d*; rACC: rostral anterior cingulate cortex; cACC: caudal ACC; PCC: posterior cingulate cortex; lOFC: lateral orbitofrontal cortex; mOFC: medial OFC. Cortisol contrast represents a sex-independent cortisol increase using the whole sample. Significant results are given in bold.

**Table S7.** Results of PALM calculations for cortical thickness of the right hemisphere.

| Contrast | rACC | cACC | PCC | Parahippocampus | lOFC | mOFC | Insula | Precuneus |
|---|---|---|---|---|---|---|---|---|
| **Cortisol** | | | | | | | | |
| ***t*** | .142 | **2.037** | 1.051 | -.953 | 1.131 | 1.202 | -.048 | -.126 |
| $p_{uncorr}$ | .886 | **.046** | .301 | .344 | .263 | .229 | .965 | .902 |
| $p_{FDR}$ | .965 | .366 | .550 | .550 | .550 | .550 | .965 | .965 |
| ***d*** | .058 | .834 | .430 | -.390 | .463 | .492 | -.020 | -.052 |
| **Male** | | | | | | | | |
| ***t*** | .791 | .849 | .854 | -.450 | 1.077 | .366 | .446 | -.336 |
| $p_{uncorr}$ | .420 | .397 | .396 | .649 | .277 | .715 | .652 | .773 |
| $p_{FDR}$ | .733 | .733 | .733 | .733 | .733 | .733 | .733 | .733 |
| ***d*** | .382 | .410 | .412 | -.217 | .519 | .177 | .215 | -.162 |
| **Female** | | | | | | | | |
| ***t*** | -.397 | 1.894 | .677 | -.848 | .615 | 1.216 | -.382 | .088 |
| $p_{uncorr}$ | .687 | .060 | .506 | .390 | .539 | .226 | .704 | .929 |
| $p_{FDR}$ | .805 | .478 | .805 | .805 | .805 | .805 | .805 | .929 |
| ***d*** | -.199 | .948 | .339 | -.425 | .308 | .609 | -.191 | .044 |
| **Male > Female** | | | | | | | | |
| ***t*** | .783 | -1.036 | -.049 | .424 | .132 | -.770 | .570 | -.267 |
| $p_{uncorr}$ | .433 | .296 | .960 | .667 | .895 | .444 | .565 | .792 |
| $p_{FDR}$ | .960 | .960 | .960 | .960 | .960 | .960 | .960 | .960 |
| ***d*** | .374 | -.495 | -.023 | .202 | .063 | -.367 | .272 | -.128 |
| **Female > Male** | | | | | | | | |
| ***t*** | -.783 | 1.036 | .049 | -.424 | -.132 | .770 | -.570 | .267 |
| $p_{uncorr}$ | .433 | .296 | .960 | .667 | .895 | .444 | .565 | .792 |
| $p_{FDR}$ | .960 | .960 | .960 | .960 | .960 | .960 | .960 | .960 |
| ***d*** | -.374 | .495 | .023 | -.202 | -.063 | .367 | -.272 | .128 |

$p_{uncorr}$: uncorrected *p*-values; $p_{FDR}$: FDR-corrected *p*-values; *d*: Cohen's *d*; rACC: rostral anterior cingulate cortex; cACC: caudal ACC; PCC: posterior cingulate cortex; lOFC: lateral orbitofrontal cortex; mOFC: medial OFC. Cortisol contrast represents a sex-independent cortisol increase using the whole sample. Significant results are given in bold.

**Table S8.** Results of PALM calculations for subcortical volume of the left hemisphere.

| Contrast | | Thalamus | Ncl. Caudatus | Ncl. Accumbens | Putamen | Hippocampus | Amygdala |
|---|---|---|---|---|---|---|---|
| **Cortisol** | | | | | | | |
| | $t$ | -.489 | 1.530 | .416 | .257 | -.137 | .332 |
| | $p_{uncorr}$ | .628 | .126 | .680 | .797 | .889 | .744 |
| | $p_{FDR}$ | .889 | .758 | .889 | .889 | .889 | .889 |
| | $d$ | -.197 | .616 | .168 | .103 | -.055 | .134 |
| **Male** | | | | | | | |
| | $t$ | -.383 | 1.926 | .273 | .384 | .254 | .122 |
| | $p_{uncorr}$ | .703 | .054 | .778 | .705 | .796 | .899 |
| | $p_{FDR}$ | .899 | .323 | .899 | .899 | .899 | .899 |
| | $d$ | -.184 | .928 | .132 | .185 | .123 | .059 |
| **Female** | | | | | | | |
| | $t$ | -.328 | .500 | .317 | .040 | -.353 | .322 |
| | $p_{uncorr}$ | .740 | .618 | .759 | .969 | .726 | .753 |
| | $p_{FDR}$ | .911 | .911 | .911 | .969 | .911 | .911 |
| | $d$ | -.163 | .248 | .157 | .020 | -.175 | .160 |
| **Male > Female** | | | | | | | |
| | $t$ | .042 | .725 | -.097 | .193 | .436 | -.191 |
| | $p_{uncorr}$ | .969 | .476 | .919 | .848 | .664 | .849 |
| | $p_{FDR}$ | .969 | .969 | .969 | .969 | .969 | .969 |
| | $d$ | .020 | .346 | -.046 | .092 | .208 | -.091 |
| **Female > Male** | | | | | | | |
| | $t$ | -.042 | -.725 | .097 | -.193 | -.436 | .191 |
| | $p_{uncorr}$ | .969 | .476 | .919 | .848 | .664 | .849 |
| | $p_{FDR}$ | .969 | .969 | .969 | .969 | .969 | .969 |
| | $d$ | -.020 | -.346 | .046 | -.092 | -.208 | .091 |

$p_{uncorr}$: uncorrected $p$-values; $p_{FDR}$: FDR-corrected $p$-values; $d$: Cohen's $d$. Cortisol contrast represents a sex-independent cortisol increase using the whole sample. Significant results are given in bold.

**Table S9.** Results of PALM calculations for subcortical volume of the right hemisphere.

| Contrast | | Thalamus | Ncl. Caudatus | Ncl. Accumbens | Putamen | Hippocampus | Amygdala |
|---|---|---|---|---|---|---|---|
| **Cortisol** | | | | | | | |
| | ***t*** | 1.023 | 1.861 | .467 | -.289 | -.440 | -.750 |
| | $p_{uncorr}$ | .300 | .065 | .652 | .768 | .655 | .448 |
| | $p_{FDR}$ | .768 | .392 | .768 | .768 | .768 | .768 |
| | ***d*** | .412 | .750 | .188 | -.117 | -.177 | -.302 |
| **Male** | | | | | | | |
| | ***t*** | .017 | **2.303** | .468 | .871 | .242 | .588 |
| | $p_{uncorr}$ | .986 | **.023** | .639 | .391 | .810 | .561 |
| | $p_{FDR}$ | .986 | .137 | .958 | .958 | .972 | .958 |
| | ***d*** | .008 | 1.110 | .225 | .420 | .116 | .283 |
| **Female** | | | | | | | |
| | ***t*** | 1.252 | .637 | .239 | -.987 | -.718 | -1.352 |
| | $p_{uncorr}$ | .203 | .527 | .807 | .323 | .465 | .175 |
| | $p_{FDR}$ | .610 | .632 | .807 | .632 | .632 | .610 |
| | ***d*** | .621 | .316 | .119 | -.490 | -.356 | -.671 |
| **Male > Female** | | | | | | | |
| | ***t*** | -1.007 | .835 | .081 | 1.312 | .725 | 1.443 |
| | $p_{uncorr}$ | .319 | .412 | .938 | .192 | .474 | .148 |
| | $p_{FDR}$ | .569 | .569 | .938 | .569 | .569 | .569 |
| | ***d*** | -.481 | .399 | .038 | .627 | .346 | .689 |
| **Female > Male** | | | | | | | |
| | ***t*** | 1.007 | -.835 | -.081 | -1.312 | -.725 | -1.443 |
| | $p_{uncorr}$ | .319 | .412 | .938 | .192 | .474 | .148 |
| | $p_{FDR}$ | .569 | .569 | .938 | .569 | .569 | .569 |
| | ***d*** | .481 | -.399 | -.038 | -.627 | -.346 | -.689 |

$p_{uncorr}$: uncorrected *p*-values; $p_{FDR}$: FDR-corrected *p*-values; *d*: Cohen's *d*. Cortisol contrast represents a sex-independent cortisol increase using the whole sample. Significant results are given in bold.

**Table S10.** Cortisol increase prediction scores using different brain measure and confound correction settings.

| Brain Measures | Metric | | Metric | |
|---|---|---|---|---|
| | *r* | $p_{permutation}$ | *EV* | $p_{permutation}$ |
| Surface area, Thickness, Volume | .124 | .039 | .004 | .028 |
| Thickness, Volume | .029 | .383 | -.006 | .685 |

*r,* Pearson's correlation coefficient between true and predicted cortisol increase values; $P_{permutation}$, *p*-values derived from permutation testing, EV, Explained Variance.

### Supplementary Figures

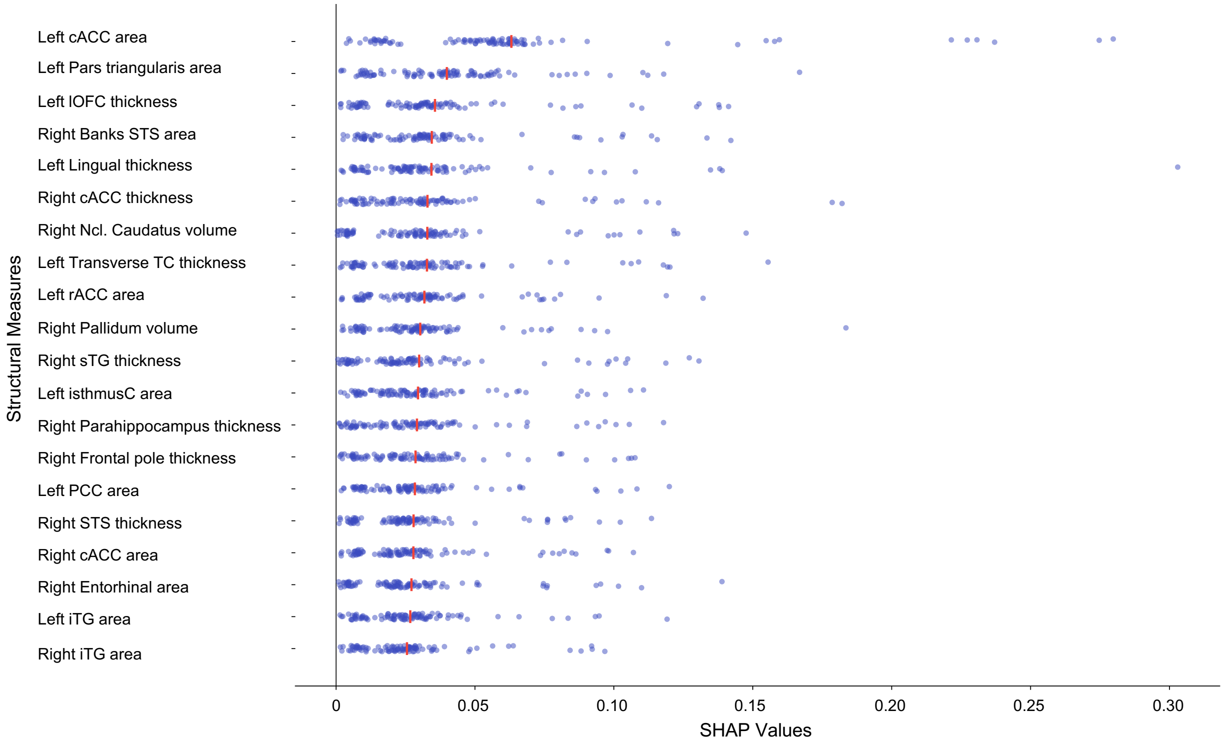


**Figure S1.** The most informative structural measures for predicting cortisol increase, as measured by SHAP values derived from a ridge regression model within each cross-validation fold. The absolute extent of feature contribution is depicted by SHAP values, with each point representing the median of the absolute SHAP values within each cross-validation fold. Mean SHAP values of each brain measure are indicated by a red line. The brain regions are denoted as follows: cACC, caudal anterior cingulate cortex; STS, superior temporal sulcus; lOFC, lateral orbitofrontal cortex; TC, temporal cortex; Ncl. Caudatus, nucleus caudatus; isthmusC, isthmus cingulate; sTG, superior temporal gyrus; PCC, posterior cingulate cortex; iTG, inferior temporal gyrus.
